# Using Attenuated Total Reflection (ATR) to Investigate the Temperature Dependent Dielectric Properties of Tympanic Membrane, Ear Canal, and Muscle tissues

**DOI:** 10.1101/2023.08.10.552862

**Authors:** Reza Shams, Zoltan Vilagosh, David Sly

## Abstract

The attenuated total reflection (ATR) setup, equipped with a diamond crystal and operating in a mixed reflection/transmission mode, demonstrated a superior and efficient capacity for investigating temperature-related interactions of biological materials at the THz-far infrared beamline at the Australian Synchrotron. This methodology was employed explicitly to investigate the temperature-driven variations in reflectance of biological tissues, such as the tympanic membrane, skeletal muscle, and brain samples, in addition to the interaction of water with THz radiation. Uniquely, the technique detected a characteristic ‘crossover flare’ feature in the spectral scan, a trait inherent to water and water-based compounds. It also identified a ‘quiet zone’ feature, a region exhibiting no temperature-dependent reflectance variation at higher frequencies. Remarkably, this approach required minimum sample preparation and was non-destructive, enabling the investigation of a range of tissue temperatures to ascertain the influence of temperature on the reflection and absorption dynamics of THz radiation.

## Introduction

The study of temperature dependent radiofrequency interactions in biological tissues has been a topic of interest in recent years. One area of focus is the use of terahertz (THz) frequencies, which lie between microwave and infrared on the electromagnetic spectrum. These frequencies have unique absorption characteristics in water and other biomolecules, making them useful for sensing and imaging applications in biomedical sciences and biological studies. In order to investigate the effect of temperature on THz absorption in various tissues, researchers have employed various techniques such as spectroscopy and imaging. The results have shown that there is a strong dependence on temperature, with absorption increasing at higher temperatures (Vilagosh et al., 2022).

Research has revealed that temperature significantly impacts the interaction of radio frequencies in the THz range with water molecules. As temperature increases, the absorption and transmission coefficients of water also increase in the THz range (Mumtaz et al., 2020; Ronne et al., 1997). This is attributed to the increased molecular motion and rotation of water molecules at higher temperatures, resulting in increased absorption of THz radiation. The changes in dielectric properties of water, such as the real and imaginary parts of the dielectric constant and loss tangent, primarily drive this interaction (Zhou et al., 2019). However, the relationship between temperature and absorption is non-linear, with the absorption coefficient showing a non-linear increase with temperature (Vilagosh et al., 2022).

This non-linear relationship can be attributed to the complex behavior of water molecules, which exhibit different types of motion at different temperatures. The rotational motion of water molecules primarily influences the absorption coefficient at lower temperatures, while the translational motion, or the movement of water molecules as a whole, becomes more significant at higher temperatures (Elgabarty et al., 2020; Śmiechowski et al., 2016). Additionally, the increased thermal energy at higher temperatures also causes water molecules to vibrate more, further increasing absorption. The combination of rotational, translational, and vibrational motion results in the non-linear increase of the absorption coefficient with temperature in the THz range (Xu & Havenith, 2015), with the coefficient leveling off at higher temperatures indicating a saturation point.

Previous findings on the temperature dependent interaction of THz radiation with water molecules have important implications for the study of biological tissues. Water is present in all biological tissues, and the absorption and transmission coefficients of THz radiation are directly influenced by the temperature of the free and bound water within the tissue (Vilagosh et al., 2022). This suggests that different tissues with varying water content and temperature may have different interactions with THz radiation. For example, the thick skin of an organism, with lower water content and lower temperature, may be well-evolved to deal with the environmental force of THz radiation. However, the thin tympanic membrane, with high water content and high temperature, may not be able to deal with this interaction as effectively. This highlights the importance of considering the temperature dependent interaction of THz radiation with water in the study of biological tissues.

Examining the temperature dependent THz interaction with materials using transmission apparatus can be challenging for a few reasons. One reason is that the THz frequency range is relatively low, which makes it difficult to accurately measure small changes in transmission. Additionally, the absorption and transmission of THz waves in a material can be highly dependent on the sample’s temperature, making it difficult to separate the effects of temperature from the effects of the material itself. Additionally, THz radiation is weakly interacting with materials, making it challenging to obtain strong enough signal to noise ratio. Furthermore, many materials are opaque in the THz range, making it difficult to obtain accurate transmission measurements. An alternative method called ATR can provide a solution for some of the difficulties mentioned above and can be more accurate, can provide a way to separate the effects of temperature from the effects of the material itself, overcome the challenges of weak signal to noise ratio and opaque materials in the THz range, and also establish clear trends and correlations between temperature and THz interactions (Ryu et al., 2021).

ATR has been demonstrated as an effective method for studying the temperature dependent interaction of THz radiation with materials. This technique utilises the evanescent wave generated at the interface between a prism and a sample to probe the sample’s absorption and transmission properties (Vilagosh et al., 2022). The sensitivity of this method to the refractive index of the sample makes it particularly useful for studying the temperature dependent dielectric properties of materials, including water, in the THz range. Studies have shown that the use of ATR allows for the measurement of temperature dependent absorption and transmission coefficients of materials, providing valuable insight into the temperature dependent behavior of THz radiation in these materials. The ability to measure temperature dependent changes in the dielectric properties of materials with ATR, make it a useful tool for studying the temperature dependent interaction of THz radiation with biological tissue.

In addition to the advantages mentioned above, ATR also offers several conveniences when it comes to examining the temperature dependent THz interactions with materials. One of the main advantages is its ability to change the temperature of the material quickly and easily, allowing for rapid investigation of the THz interactions at different temperatures. This is particularly useful for materials that have a limited temperature range or are sensitive to temperature changes. Additionally, ATR measurements are typically faster and easier to perform than transmission measurements, making it a more efficient method for investigating the temperature dependent THz interactions with materials. This makes it an ideal technique for researchers who need to perform multiple measurements in a short amount of time. In summary, ATR is a more convenient, fast, and easy to use alternative for examining the temperature dependent THz interactions with materials.

To investigate the interaction of THz with materials using ATR, it is important to account for both reflection and absorption. The reflectivity of a material is determined by its refractive index and the angle of incidence of the THz radiation. The refractive index of a material is directly related to its dielectric properties, and it is this relationship that determines the amount of THz radiation that is reflected by a material (Ryu et al., 2021; Vilagosh et al., 2022). As the angle of incidence increases, the reflectivity of the material decreases. This is due to the increased absorption of THz radiation as it penetrates deeper into the material. Additionally, the thickness and composition of a material also affect its reflective properties in the THz range. To fully understand the interaction of THz with a material, it is important to consider both the reflective and absorptive properties of the material.

It is hypothesized that the interaction of THz radiation with biological tissue is heavily dependent on the presence and amount of water within the tissue. As water has a higher absorption coefficient in the THz range, tissues with higher water content are expected to have a greater absorption of THz radiation. Additionally, it is also hypothesized that temperature plays a crucial role in the interaction of THz radiation with biological tissue. As the temperature of water increases, its absorption and transmission coefficients also increase in the THz range. Therefore, it is expected that tissues with higher temperatures will have a greater absorption of THz radiation. This hypothesis can be tested by studying the effect of water content and temperature on the absorption of THz radiation in various biological tissues.

## Method

Fresh biological tissues, including fish muscle, porcine adipose tissue, and Cane toad (Rhinella Marina) tympanic membranes were used for this study. The toad tympani were dissected and prepared at Swinburne University of Technology Health Science lab on the day of analysis. Small pieces of tissue (approximately 1 cm x 1 cm) were cut and placed onto a diamond crystal prism. The diamond ATR prism was then mounted onto the sample holder of the Australian Synchrotron Far IR Beamline for analysis.

The effect of temperature on the transmission properties of materials was investigated by acquiring spectra at temperatures between 14°C and 44°C. Spectra were acquired using the Australian Synchrotron Far IR Beamline, which is a synchrotron radiation source that provides a continuous and consistent THz beam. The beamline was equipped with a Bruker Vertex 80v Fourier transform infrared spectrometer, which allowed for spectral acquisition in the range of 20 to 800 cm^-1^. The temperature of the sample holder was controlled using a water circulator, and the temperature of the samples was monitored using a thermocouple. Spectra were acquired for both air and the tissue samples at each temperature.

### Synchrotron Far IR Beamline

The THz spectra were processed and analyzed using OPUS (2021, Version 8.0.19) software. To account for temperature-dependent variation in the air reflection output, the tissue spectral data were normalized to the air spectrum at the same temperature as the sample. The regions of the THz spectrum that exhibited poor contrast and instrument-related interferences were excluded from the analysis. To assess measurement error, repeat spectral scans were conducted at the same temperature. The detector error was estimated as ±5% in the 20 to 30 cm^-1^ region, reducing to ±0.2% in the 50 to 100 cm^-1^ region, and increasing to ±1.5% in the 500 to 700 cm^-1^ region.

Descriptive statistics were calculated for the spectral data obtained from the Australian Synchrotron Far IR Beamline and ATR measurements. The means and standard deviations of the spectral data were calculated for each tissue sample at each temperature.

## Results

Temperature has been shown to have a significant effect on the reflectance of THz radiation. An investigation was conducted by acquiring spectra with increasing temperatures (between 14°C and 44°C) using an open stage configuration and a THz-TDS system. The results indicated a temperature-dependent change in the refractive index of air, leading to differences in the amount of THz radiation reflected by the material. To account for this variation, tissue spectral data were normalized to the air spectrum at the same temperature as the sample. Certain regions of the THz spectrum were excluded from data analysis due to poor contrast, high signal to noise ratio, and erratic normalized reflectance values.

**Figure 1.**
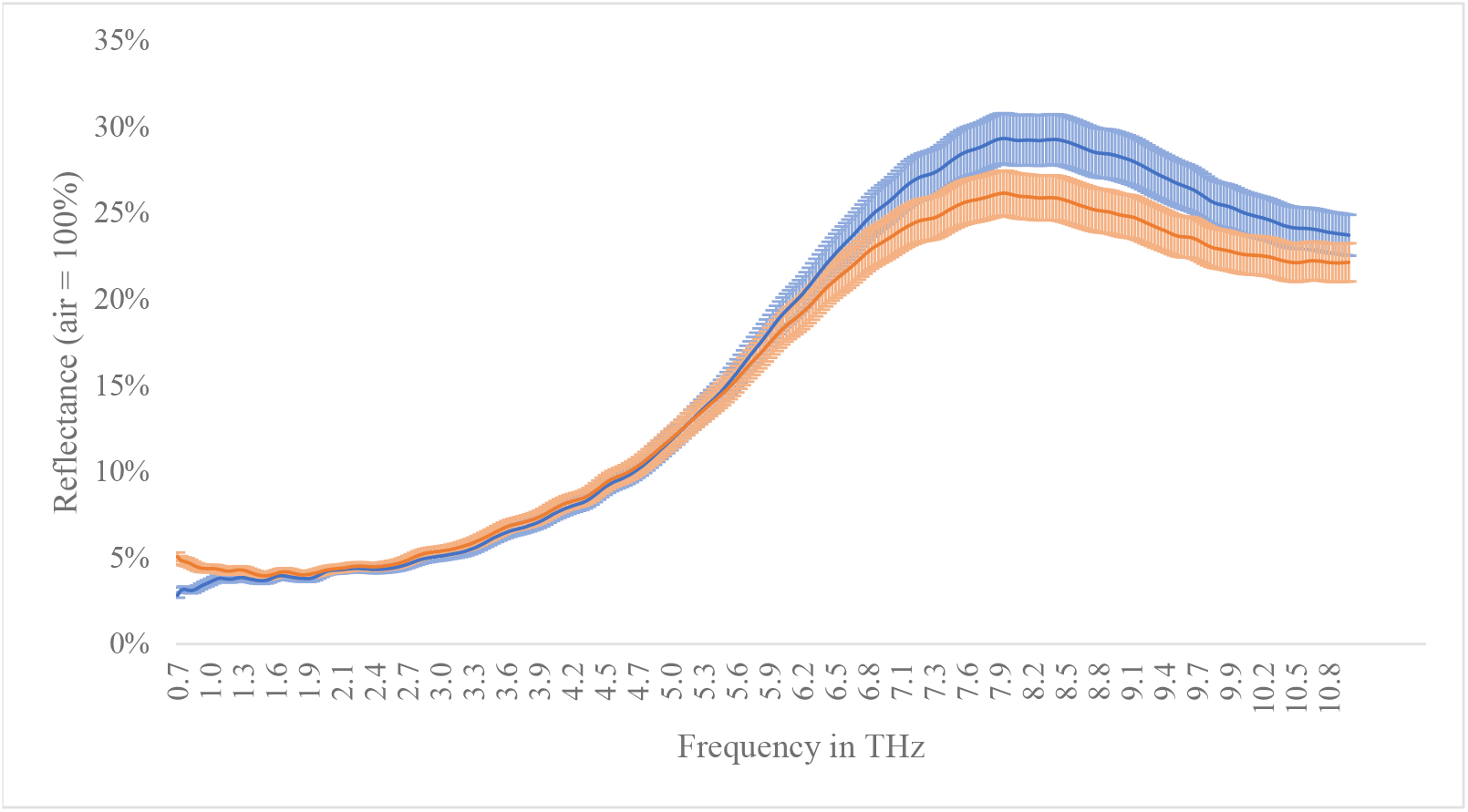
The Effect of Temperature Variation on ATR Spectra at 14 °C and 44 °C on Water Sample *Note*. The variation between the 14 °C (blue line) and 44 °C (orange line) spectra was not uniform, varying by 0.5% to 5%, depending on the frequency.

To account for this temperature-dependent variation in the air reflection output, the tissue spectral data were normalized to the air spectrum at the same temperature as the sample. In other words, spectra acquired at 14 °C were normalized to the 14 °C air spectrum, and spectra acquired at 44 °C were normalized to the air spectrum acquired at 44 °C. This normalization step is important as it allows for a more accurate comparison of the THz transmission properties of the tissue samples at different temperatures.

Based on the results of the temperature variation study within the apparatus, certain regions of the THz spectrum were excluded from the data analysis. The region below 0.7 THz was omitted due to poor contrast, high signal to noise ratio and erratic normalized reflectance values. Despite the low contrast, the data in the 0.7 THz to 2.0 THz range was included as it is within the range interest on THz application in future technologies. However, the data in the region above 10.0 THz was disregarded as it exhibited an instrument-related interferences.

To assess the measurement error, repeat spectral scans were conducted at the same temperature. The error was estimated to be ±5% in the 0.7 THz to 1.0 THz region, reducing to ±0.2% in the 3.0 THz to 6.0 THz region, and increasing to ±1.5% in the 9.0 THz to 10.0 THz region. This indicates that the measurement error is relatively low in the middle of the THz spectrum, but increases at the higher and lower frequency regions.

The data analysis of pure water and biological tissue samples revealed a similar pattern of reflection, primarily influenced by the refractive index (n) at frequencies below 5.0 THz, and by the absorption coefficient (α) at frequencies above 5.0 THz. The “quiet point,” where the influence of n and α balanced out, was determined to be at 5.2 THz. Fish muscle showed a higher reflectance compared to other tissues in the < 4 THz range, but the “quiet point” was located in the same region as the other tissues.

**Figure 2.**
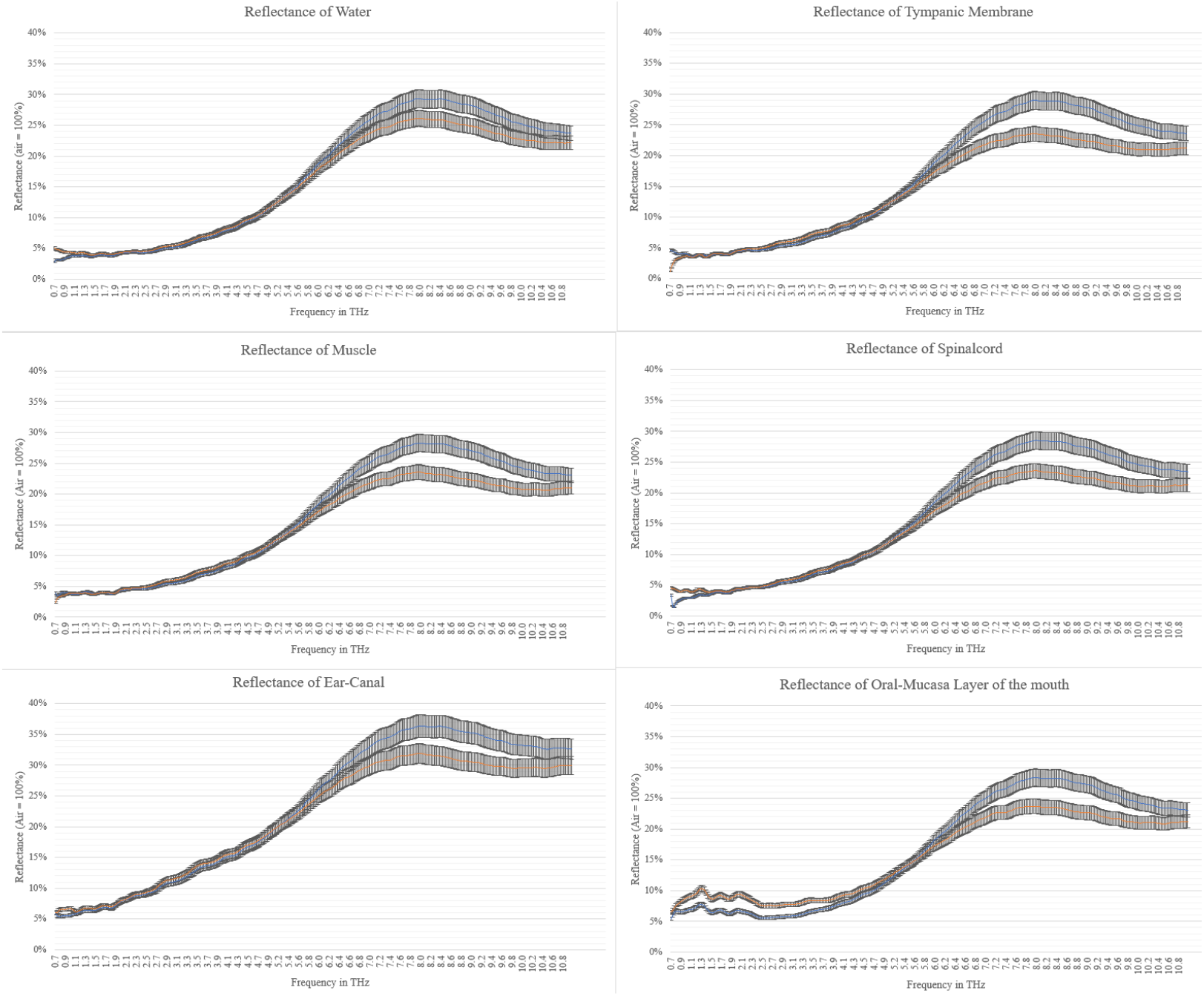
The Effect of Temperature Variation on ATR Spectra at 14 °C and 44 °C with Water and Biological Tissue in Place. *Note*. The variation between the 14 °C and 44 °C spectra was not uniform, varying by 0.5% to 5%, depending on the frequency.

The reflectance spectrum of porcine oral mucosa layer tissue highlighted a distinct temperature-dependent “crossover flare” and “quiet point,” with the “quiet point” occurring at 3.5 THz. This suggests that the refractive index of the tissue is approximately 1.68 at this frequency. At frequencies below 2.0 THz, the temperature-dependent spectral signature of the tissue closely resembled that of fat, with similar variations in reflectivity.

**Figure 3.**
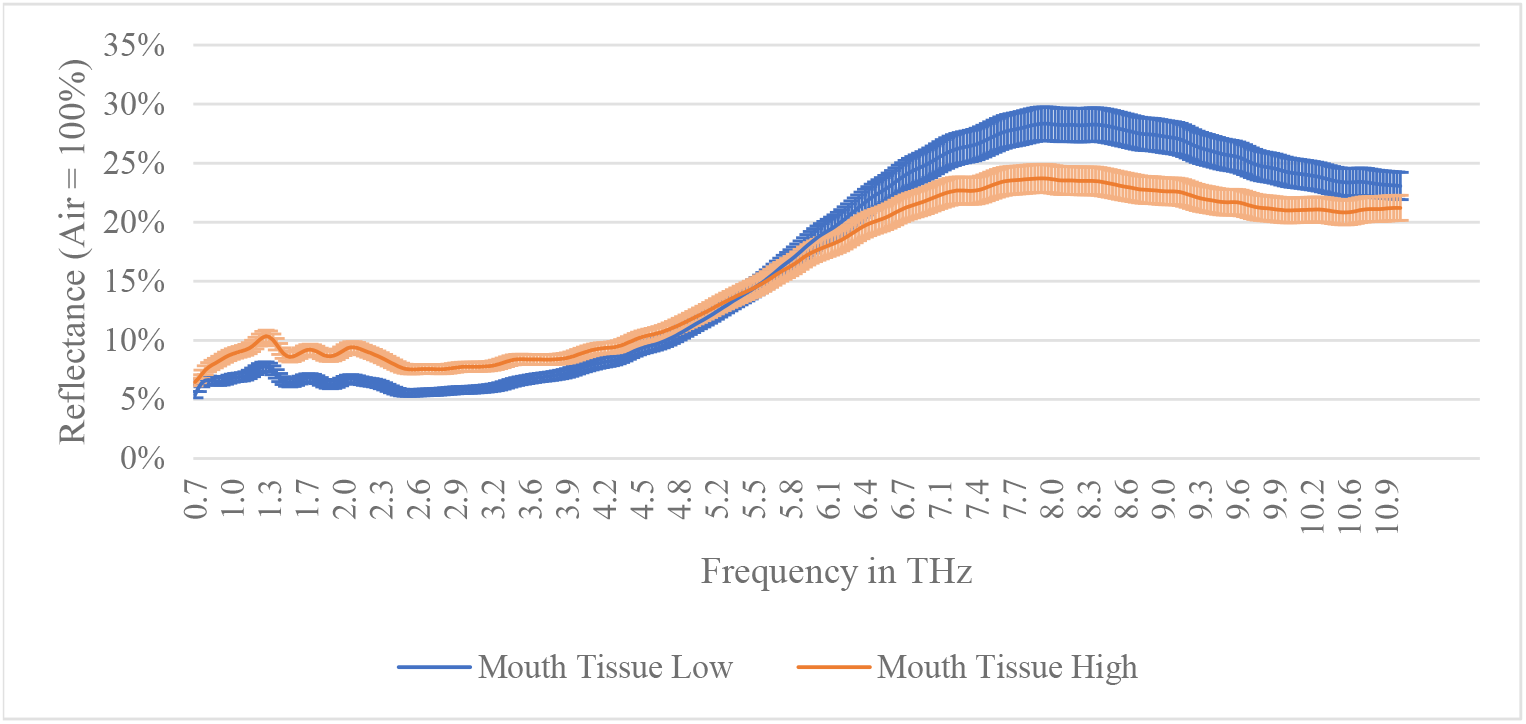
The Effect of Temperature Variation on ATR Spectra at 14 °C and 44 °C with Porcine Oral Mucosa Layer Tissue in Place. *Note*. The variation between the 14 °C and 44 °C spectra was not uniform, varying by 0.5% to 5%, depending on the frequency.

## Discussion

This study investigated the effect of temperature and water content on the reflectance properties of biological tissues in the THz frequency range. The results showed that temperature variation within the apparatus has a significant effect on the reflectance properties of THz radiation, likely due to temperature-dependent changes in the refractive index of air. To account for this variation, the tissue spectral data were normalized to the air spectrum at the same temperature as the sample. Certain regions of the THz spectrum were excluded from the data analysis due to poor contrast, high signal to noise ratio, and erratic normalized reflectance values.

The study also demonstrated that water content plays a crucial role in the reflectance properties of biological tissues in the THz frequency range. The spectral signature of the tissue below 2.0 THz was observed to be similar to that of adipose tissues, with similar variations in reflectivity. The high-water content of both the tissue and the fat may contribute to the similarities observed in their THz absorption and reflection properties. The data also revealed a similar pattern of reflection in pure water and biological tissues, where the temperature-dependent changes in reflection were primarily influenced by the refractive index (n) at frequencies below 5.0 THz and by the absorption coefficient (α) at frequencies above 5.0 THz. The “quiet point,” where the influence of n and α balanced out, was determined to be at 5.2 THz.

In addition to demonstrating the roles of temperature and water content in THz reflectance properties, the study identified important implications for future THz applications. The reflectance spectrum of porcine oral mucosa layer tissue highlighted a distinct temperature-dependent “crossover flare” and “quiet point,” with the “quiet point” occurring at 3.5 THz. This suggests that the refractive index of the tissue is approximately 1.68 at this frequency, which can be useful for the detection of early-stage oral cancer. Furthermore, the findings underscore the need for a better understanding of the role of water and temperature in THz characterization of materials, which could lead to the development of more accurate and reliable THz techniques for the characterization of biological tissues and other materials with high water content.

Crucially, this study reveals that the interaction of THz radiation with the tympanic membrane follows a similar pattern to that of water, emphasizing the significant role water absorption plays in the interaction between tissue and electromagnetic radiation. This suggests that the high-water content of both the tissue and fat might contribute to the observed similarities in their THz absorption and reflection properties. Additionally, the results highlight that at higher frequencies, specifically above 5.0 THz, temperature can substantially affect and influence the absorption and hence, the subsequent potential thermal effect. In particular, temperature-dependent changes in reflection were found to be largely influenced by the refractive index at frequencies below 5.0 THz and by the absorption coefficient at frequencies above 5.0 THz.

The findings from this study have crucial implications, most notably suggesting a differential response of various body tissues to THz radiation. In contrast to the skin, which is substantially influenced by ambient temperatures that are relatively lower than the body’s core temperature, the interaction of the tympanic membrane with THz radiation appears to be moderated and indeed enhanced. More specifically, as the temperature elevates, there is a notable increase in the normalized reflectance, ranging between 2% to 5%, compared to lower temperature conditions. This significant increase in reflectance at higher temperatures implies that the tympanic membrane, typically surrounded by a temperature of 37 degrees Celsius, may exhibit increased absorption of THz radiation.

It is important to note that the results presented in this study are limited to porcine and toad tissues and may not necessarily generalise to human tissues. Future research is needed to investigate the reflectance properties of other human tissues in the THz frequency range and to explore the relationship between reflectance properties and structural or morphological properties of the tissue. Nonetheless, the findings of this study provide important insights into the reflectance properties of biological tissues in the THz frequency range and have significant implications for future THz applications and characterization of materials.

This study underscores the significant roles that temperature and water content play in the interaction of THz radiation with biological tissues, including the tympanic membrane and muscular tissues. These tissues, known for their high water content, present similar patterns of THz absorption and reflection. It becomes particularly striking to observe the increased influence of temperature on absorption at frequencies exceeding 5.0 THz across all samples. An intriguing ‘quiet point’ was identified, where the impacts of refractive index and absorption coefficient reach equilibrium, which holds noteworthy implications for a comprehensive spectral understanding of biological tissues. Collectively, these insights stress the importance of a more profound comprehension of the interplay between water, temperature, and THz characterization of biological materials.

## Notes

### Competing Interest Statement

The authors have declared no competing interest.

## Reference

Elgabarty, H., Kampfrath, T., Bonthuis, D. J., Balos, V., Kaliannan, N. K., Loche, P., … & Sajadi, M. (2020). Energy transfer within the hydrogen bonding network of water following resonant terahertz excitation. Science Advances, 6(17), eaay7074. https://doi.org/10.1126/sciadv.aay7074

Ro/nne, C., Thrane, L., Åstrand, P. O., Wallqvist, A., Mikkelsen, K. V., & Keiding, S. R. R. (1997). Investigation of the temperature dependence of dielectric relaxation in liquid water by THz reflection spectroscopy and molecular dynamics simulation. The Journal of chemical physics, 107(14), 5319–5331. https://doi.org/10.1063/1.474242

Ryu, M., Ng, S. H., Anand, V., Lundgaard, S., Hu, J., Katkus, T., … & Morikawa, J. (2021). Attenuated total reflection at THz wavelengths: Prospective use of total internal reflection and polariscopy. Applied Sciences, 11(16), 7632. https://doi.org/10.3390/app11167632

Śmiechowski, M., Schran, C., Forbert, H., & Marx, D. (2016). Correlated particle motion and THz spectral response of supercritical water. Physical Review Letters, 116(2), 027801. https://doi.org/10.1103/PhysRevLett.116.027801

Vilagosh, Z., Appadoo, D., Foroughimehr, N., Shams, R., Sly, D., Juodkazis, S., … & Wood, A. W. (2022). Tissue Characterization Using Synchrotron Radiation at 0.7 THz to 10.0 THz with Extended ATR Apparatus Techniques. Sensors, 22(21), 8363. https://doi.org/10.3390/s22218363

Xu, Y., & Havenith, M. (2015). Perspective: Watching low-frequency vibrations of water in biomolecular recognition by THz spectroscopy. The Journal of chemical physics, 143(17). https://doi.org/10.1063/1.4934504

Zhou, J., Rao, X., Liu, X., Li, T., Zhou, L., Zheng, Y., & Zhu, Z. (2019). Temperature dependent optical and dielectric properties of liquid water studied by terahertz time-domain spectroscopy. AIP Advances, 9(3). https://doi.org/10.1063/1.5082841

